# Privacy-Preserving Statistical Analysis of Genomic Data using Compressive Mechanism with Haar Wavelet Transform*

**DOI:** 10.1101/2022.04.10.487816

**Authors:** Akito Yamamoto, Tetsuo Shibuya

## Abstract

To promote the use of personal genome information in medicine, it is important to analyze the relationship between diseases and the human genomes. Therefore, statistical analysis using genomic data is often conducted, but there is a privacy concern with respect to releasing the statistics as they are. Existing methods to address this problem using the concept of differential privacy cannot provide accurate outputs under strong privacy guarantees, making them less practical. In this study, for the first time we investigate the application of a compressive mechanism to genomic statistical data and propose two approaches. The first is to apply the normal compressive mechanism to the statistics vector along with an algorithm to determine the number of nonzero entries in a sparse representation. The second is to alter the mechanism based on the data, aiming to release significant SNPs with a high probability. In this algorithm, we apply the compressive mechanism with the input as a sparse vector for significant data and the Laplace mechanism for non-significant data. Using the Haar transform for the wavelet matrix in the compressive mechanism is advantageous to determine the number of nonzero elements and the amount of noise. In addition, we theoretically prove that our proposed method achieves *ϵ*-differential privacy. We evaluated our methods in terms of accuracy, rank error, and run time compared to the Laplace and exponential mechanisms. The results show that our second method in particular can guarantee both high privacy assurance as well as utility. The Python implementation of our experiments is available at https://github.com/ay0408/CompLaplace.

## 1 Introduction

In recent years, the availability of the information on human genome has increased, and more opportunities to utilize these data for personalized medicine are expected [20]. Therefore, it is essential to analyze the relationship between diseases and genomic information, and investigations, such as genome-wide association studies (GWAS) are often conducted. However, these tests contain sensitive personal information, and there is a risk of identifying an individual [33]. In fact, several studies [21, 30, 34] have been conducted on methods to identify individuals by using large-scale genomic data. As a result, the NIH has stopped releasing aggregate GWAS data [40], making it difficult to freely utilize these data.

Therefore, it is desirable to develop techniques for secure genomic analysis and methods for the private publication of statistical data. Various studies have been conducted on protecting data privacy using cryptography, secret sharing, and multiparty computation [6, 7, 11, 41], but they need additional protections of calculation results. Thus, in this study, we focus on the concept of differential privacy [15], a strategy that is being considered for protecting and utilizing genomic data [5, 17, 32, 38]. The concept of differential privacy is based on the fact that when some degree of perturbation is introduced in the original dataset, it becomes almost impossible to determine whether the dataset contains information pertaining to a specific individual. This concept might enable privacy protection even if noisy data are released to the public.

However, existing methods [5, 17, 36, 38] that employ the strategy of differential privacy have the limitation that there is a lot of differences between noisy data and the original data. These methods require a large budget *ϵ* for ensuring the utility of the data while achieving high privacy. The exponential mechanism using the shortest Hamming distance scores [23] provides relatively high accuracy, but it requires the calculation of the score for each statistic every time, and few efficient algorithms exist for this purpose. To circumvent this problem, methods [22, 31] employing probabilistic modeling have been proposed, but there is still room for improvement in terms of accuracy.

In this study, we propose new methods providing differentially privacy with a high utility to release genomic data. The goal of this study is to privately release the top *K* significant single nucleotide polymorphisms (SNPs). First, we consider subjecting genomic data to a compressive mechanism [25], which is based on compressed sensing and is suitable for vector data with a sparse representation. It has been studied for publication of histograms [2] and for use in network systems [12]; however, to the best of our knowledge, this study is the first to apply this method to genomic statistical data. We also present an algorithm to determine the number of nonzero entries in a sparse vector obtained using compressed sensing to make the compressive mechanism more reliable. However, because statistical data do not always have a sparse distribution, we consider the input data as a sparse vector and proposed another mechanism that combines the compressive mechanism and Laplace mechanism [16]. This mechanism is aimed at enabling the release of significant SNPs with a high probability by distinguishing between significant and non-significant data. In our methods, we use the Haar wavelet transform [18] as an orthonormal basis in the compressive mechanism as it is easy to determine the number of nonzero entries because the inverse matrix matches the transpose matrix, and as the *sensitivity* can be analyzed easily to calculate the amount of noise. We also theoretically guarantee that our methods achieve *ϵ*-differential privacy.

In our experiments, we compare the results obtained using our two methods with those obtained using the Laplace mechanism [16] and the exponential mechanism [27], the key mechanisms for ensuring differential privacy. First, we evaluate the accuracy and rank error of the top K outputs, and show that our methods have high utility. We also measure the run time while varying the size of the datasets. In addition, we subject real data [24] to our methods to verify their usefulness.

In Section 2, we describe the basic and preliminary assumptions. In Section 3, we present *ϵ*-differentially private methods using the compressive mechanism with the Haar wavelet transform to release the top *K* significant SNPs based on genomic statistics data. In Section 4, we evaluate their utility based on simulations and real data. In Section 5, we summarize our study and directions for future work.

## 2 Preliminaries

### 2.1 Differential Privacy

Differential privacy [15] is a framework developed in the field of cryptography to protect personal data; currently, it is being extensively considered for releasing genomic data [17, 32, 38]. It is also being extensively used in the field of deep learning [1] and healthcare [3]. The concept of differential privacy is based on the idea that it is almost impossible to distinguish between two *neighboring* datasets that differ with respect to only one element. This element is determined based on the data to be protected, which often represents a single individual or one family in the case of genomic data. The privacy level in differential privacy is evaluated by the parameter *ϵ* > 0, and the following is the definition of *ϵ*-differential privacy.

#### Definition 1.

(*ϵ-Differential Privacy*)

*A randomized mechanism M is ϵ-differentially private if, for all neighboring datasets D and D′ and any S*

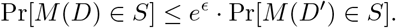

The smaller the value of *ϵ*, the stronger the privacy guarantee achieved, but the utility of such data is low. In general, the value of *ϵ* is set in the range of 0.01 to 10 [19], but when a higher privacy is required such as in the case of genomic data, it is desirable to adopt a value less than 1.

#### 2.1.1 Laplace Mechanism

The simplest mechanism to achieve *ϵ*-differential privacy is the Laplace mechanism [16], which introduces perturbations according to the *sensitivity* of the function to obtain the output. The definition of the *sensitivity* is as follows, and in this study, we view a statistic as a function.

#### Definition 2.

(*Sensitivity for the Laplace mechanism*)

*The sensitivity of a function* 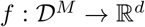 *is*

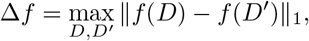

*where D*, 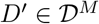 *are neighboring datasets.*

For a statistic *f* (*D*) obtained from the original dataset *D*, release of *f* (*D*) + *b* satisfies *ϵ*-differential privacy when *b* is random noise derived from a Laplace distribution with mean 0 and scale 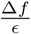 [16].

#### 2.1.2 Exponential Mechanism

The exponential mechanism [27] is a technique for providing the desired data with a high probability and has often been used to release statistically significant data [32, 38]. The desirability of the output is indicated by a score function, and we consider an exponential distribution using the *sensitivity* of the function. As a score function, the χ^2^-statistics, p-values, and shortest Hamming distance (SHD) score [23] are commonly used. The SHD score provides higher accuracy than statistics [32, 38], but for most tests, efficient methods to obtain the score has not been proposed and is not yet generally available. Therefore, in our experiments, we compare our proposed methods with the exponential mechanism using χ^2^-statistics to confirm their usefulness. The *sensitivity* for the exponential mechanism is defined as follows.

#### Definition 3.

(*Sensitivity for the exponential mechanism)*

*The sensitivity of a score function* 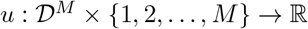 *is*

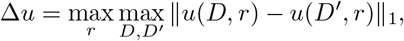

*where r* ∈ {1, 2,…, *M*} *and D*, 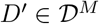 *are neighboring datasets.*

Following the above definition, we choose mechanism 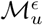, which has the distribution:

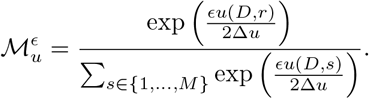

Then, releasing 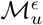 satisfies the definition of *ϵ*-differential privacy.

### 2.2 Compressive Mechanism

The compressive mechanism is a differential privacy method proposed by Li *et al.* [25]. This mechanism uses the technique of compressed sensing and has the advantage of requiring less noise than other mechanisms when the input has a sparse representation. Firstly, we briefly review the procedure involved in compressed sensing.

#### 2.2.1 Compressed Sensing

Compressed sensing was introduced by Candès *et al.* [8, 9] and Donoho [14], and has been employed in various fields, such as image processing [39] and computational biology [10].

In compressed sensing, we consider an input vector *d* ∈ ℝ^*n*^ with a sparse representation. Here, *d* can be expressed as *d* = Ψ*x*, where Ψ ∈ ℝ^*n*×*n*^ is an orthonormal basis and *x* ∈ ℝ^*n*^ is a sparse vector. Note that if *x* has *s* (< *n*) nonzero entries, then *x* is an *s*-sparse vector. The goal of compressed sensing is to recover this sparse vector *x* form the input vector *d*. The procedure is divided into the following processes: “sampling process” and “reconstruction process”, and we explain each in the following.

The sampling process reduces the size of the input from *n* to *k* (< *n*) by mapping from the input vector d to a vector *y* ∈ ℝ^*k*^. In this process, we use a random matrix Φ ∈ ℝ^*k*×*n*^ and calculate *y* = Φ*d*. For the compressive mechanism, the random matrix Φ is formed by sampling independent and identically distributed (i.i.d.) entries from a symmetric Bernoulli distribution. Half of the Φ entries are expected to be 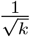 and the other half 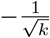, or more precisely, the following equation holds: 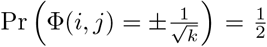. Here, when we set *k* = Θ (slog(*n/s*)), it is known that for any orthonormal basis Ψ, *A* = ΦΨ ∈ ℝ^*k*×*n*^ satisfies the restricted isometry property (RIP), a property favorable for reconstructing *x,* with a very high probability [4].

The reconstruction process calculates a vector *x* ∈ ℝ^*n*^ that satisfies *y* = ΦΨ*x*. Using *l*_1_-norm minimization, such as orthogonal matching pursuit (OMP) [28] or subspace pursuit [13], we solve the following optimization problem:

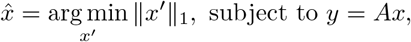

where *A* = ΦΨ. In this study, we adopt the Haar wavelet transform as Ψ.

In the compressive mechanism, we first calculate *y* = Φ*d* from the original data *d,* and then subject the vector *y* to the Laplace mechanism. Here, if the *sensitivity* of *d* is Δ_*d*_, that of *y* is expressed as 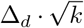 [25]. Therefore, we can introduce Laplace noise with scale 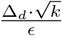 to the vector *y*. Based on the resulting *ŷ*, we can estimate a sparse vector 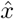 using the reconstruction process, and by computing 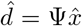, we can release privacy-protected data.

### 2.3 Haar Wavelet Transform

The Haar wavelet transform [18] is one of the simplest wavelet transforms and can be used for frequency analysis. Similar to compressed sensing, it has been employed in various fields such as image processing [29] and pattern recognition [26]. Owing to the fact that the *sensitivity* of the Haar wavelet transform is easy to analyze, it can be used along with the concept of differential privacy [35].

For the Haar wavelet transform, we use the Haar transform matrix *H* ∈ ϵ^*n*×*n*^, where *n* is a power of 2. This matrix *H* is an orthonormal basis, and the elements of H can be obtained using the following equation:

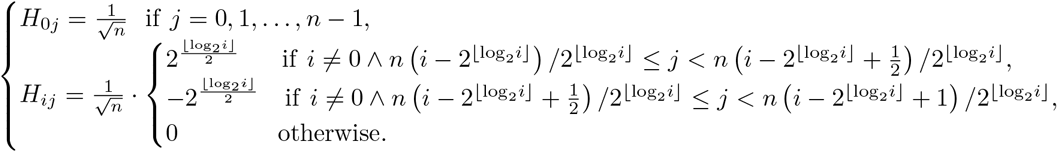

In this study, we calculate the *sensitivity* of the Haar wavelet transform and combine it with the compressive mechanism to propose a new method for releasing differentially private genomic data.

## 3 Methods

The goal of this study is to release the top *K* significant SNPs based on χ^2^-statistics in high-throughput analyses of genomic datasets, such as GWAS, while achieving *ϵ*-differential privacy. Therefore, we assume that the statistics vector *g* in the following algorithms consists of χ^2^-statistics. We first consider subjecting the genomic data to the simple compressive mechanism proposed by Liu *et al.* [25]. In this method, we focus on determining the number of nonzero entries in the sparse vector. In addition, we combine the compressive mechanism with the Haar wavelet transform and the Laplace mechanism, which can release significant SNPs with high accuracy. We also theoretically prove that this method satisfies *ϵ*-differential privacy.

### 3.1 Compressive Mechanism

In this study, we use the value of *sensitivity* according to each statistic. Several studies [17, 36, 38, 32, 37] have shown the *sensitivity* for the χ^2^-test, the Cochran-Armitage trend test, and the transmission disequilibrium test. In the future, it is expected that other statistics will also be analyzed and when we know the *sensitivity* of statistics that we want to release in the same way as in previous studies, the statistical data vector can be subjected to the compressive mechanism. Algorithm 1 represents the detailed procedure for releasing the top *K* data based on the statistics. Note that we need to introduce 2*K* times more Laplace noise [5] in each statistic than in the case where we release a single statistic. In addition, the size *k* of the vector *y* is calculated by *k* = ⌊slog(*n/s*)⌋ for ease of reconstruction.

**Algorithm 1.**
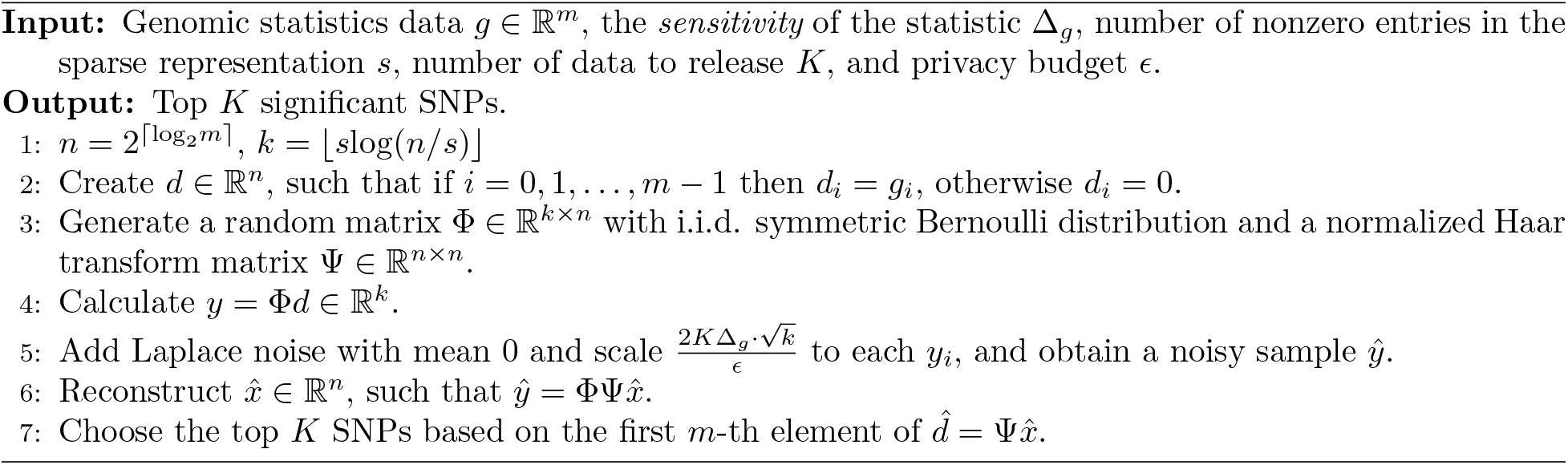
*ϵ*-differentially private algorithm for releasing the top *K* significant SNPs using the compressive mechanism.

In Algorithm 1, we assume that the number of nonzero entries of the sparse vector *x* is known. However, because the actual statistic vector may not have a sparse representation, it is necessary to establish an appropriate *s*. Therefore, we also propose an algorithm to determine the value *s.* In this algorithm, we first find *x*\* that satisfies *d* = Ψ*x*\*. Here, Ψ is the normalized Haar transform matrix, and Ψ^T^ = Ψ^-1^. Hence, *x*\* can be calculated as *x*\* = Ψ^T^*d*. Then, we describe *s* as the number of elements of *x*\* that are greater than the threshold *η.* The above procedure is shown in Algorithm 2.

**Algorithm 2.**
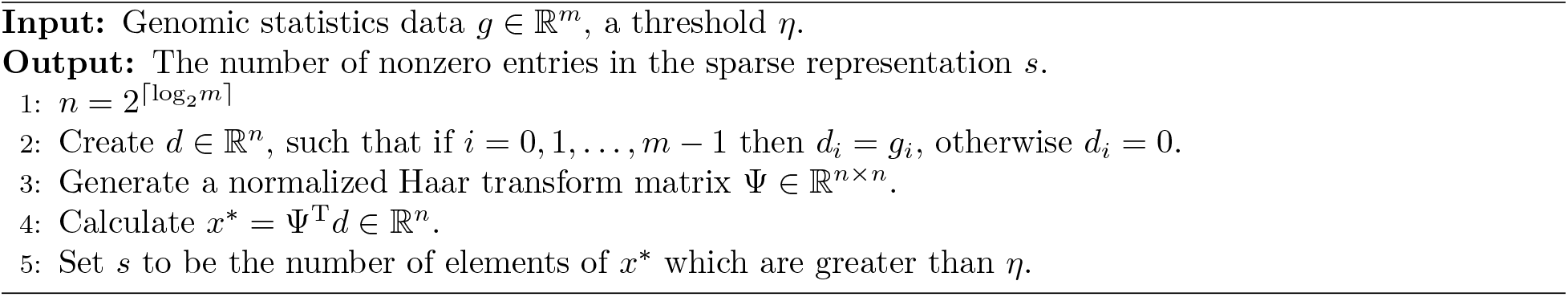
Algorithm for determining the number of nonzero entries in a sparse representation of *g* ∈ ℝ^*m*^ for Algorithm 1.

### 3.2 Compressive Mechanism + Laplace Mechanism

Algorithm 1 assumes that the input vector *g* has a sparse representation. However, this assumption does not necessarily hold, in which case the introduced noise upon using this method would be greater than that associated with the Laplace mechanism. To overcome this problem, we consider finding the vector *d* in the compressive mechanism from a sparse input vector *x*. Specifically, we first create a sparse vector *x* ∈ ℝ^*n*^ with the top 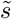 entries of the statistics vector *g* ∈ ℝ^*m*^ as nonzero elements. We then reconstruct a noisy sparse vector *x* using a methodology similar to that employed with the compressive mechanism. For the other entries, we employed the Laplace mechanism, and finally, we select the top *K* elements from all the obtained noisy statistics. The above procedure is shown in Algorithm 3.

The value 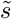 in Algorithm 3 assumes the number of high statistics in the input statistics. In general, significant statistics are often quite larger than non-significant statistics in genetic statistical tests, as shown in Manhattan plots. Because the general Laplace mechanism cannot capture this feature, we intend to differentiate the significant SNPs from non-significant SNPs by changing the mechanism employed. In addition, the amount of noise in Algorithm 3 is different from that generated upon using Algorithm 1 because the input vector in the compressive mechanism is a sparse vector *x*. We show the *sensitivity* of the Haar wavelet transform by Theorem 1.

**Algorithm 3.**
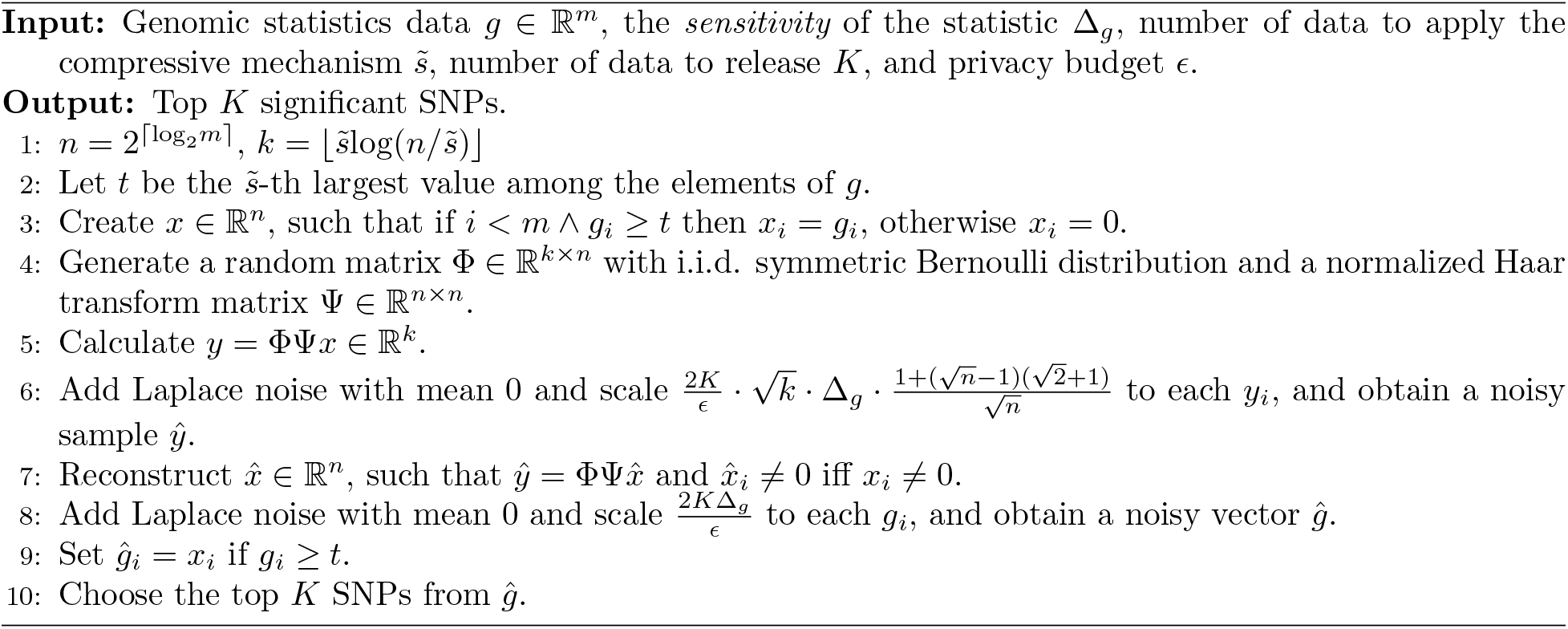
*ϵ*-differentially private algorithm for releasing the top *K* significant SNPs using the compressive mechanism with Haar wavelet transform and the Laplace mechanism.

#### Theorem 1.

*When the sensitivity of a vector x* ∈ ℝ^*n*^ *is* Δ_*x*_, *the sensitivity of the Haar wavelet transform for x is* 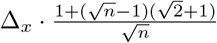.

*Proof.* The Haar wavelet transform can be defined as the following function *h* : ℝ^*n*^ → ℝ^*n*^, *h*(*x*) = Ψ*x*, where Ψ is a normalized Haar transform matrix.

When an element *x_i_* changes by *t_i_*, || *h*(*x*) ||_1_ changes by at most

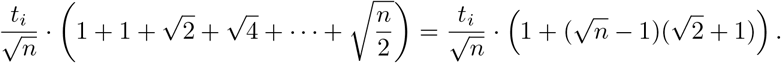

Here, the maximum change in ||*x*||_1_ is Δ_*x*_, and 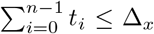 is satisfied. Therefore, the *sensitivity* of the Haar wavelet transform is 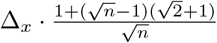.

From Theorem 1, when the *sensitivity* of the statistic is Δ_*g*_, that of *d* = Ψ*x* ∈ ℝ^*n*^ can be expressed as 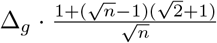. Therefore, the *sensitivity* of *y* = Φ*d* ∈ ℝ^*k*^ is 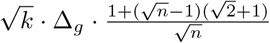, in accordance with Li *et al.* [25]. The amount of noise in the Laplace mechanism is in agreement with that proposed by Yu *et al.* [38]. In addition, we theoretically prove that Algorithm 3 satisfies *ϵ*-differential privacy by Theorem 2.

#### Theorem 2.

*Algorithm 3 satisfies *ϵ*-differential privacy.*

*Proof.* Let *U* be the set of data contained in the input vector and *W* be the set of K data obtained by Algorithm 3. In addition, let *g, g′* ∈ ℝ^*m*^ be a neighboring statistics dataset. We set 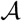 as the mechanism represented by Algorithm 3, and we will show 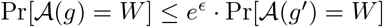.

Here, suppose that the input vector is 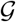, and let the noisy statistic for each element 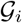 be 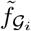 and the noisy vector *ŷ* in the compressive mechanism be 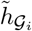.

Among the *K* output data, we assume that *p* data are chosen from the 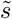 data which is subjected to the compressive mechanism, which we denote as 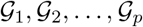, and the other data as 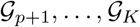.

Then we let *v*_1_, *v*_2_, …, *v_K_* ∈ ℝ be the noisy statistic of 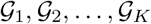, and let *z*_1_, *z*_2_, …, *z_p_* ∈ ℝ^*k*^ be the vectors for obtaining *v*_1_, *v*_2_, …, *v_p_* by the reconstruction process.

Under the above conditions, the following equation holds:

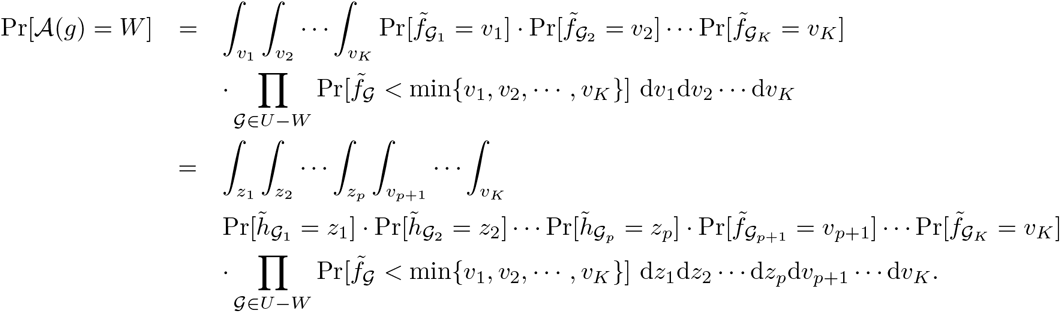

Let the sensitivity of 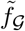 and 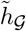 be Δ_*f*_ and Δ_*h*_, respectively. In addition, let the scale of the Laplace noise added by the Laplace mechanism and compressive mechanism be λ_*f*_ and λ_*h*_, respectively. Then, in the same way as in the discussion of Bhasker *et al.*’s Theorem 4 [5],

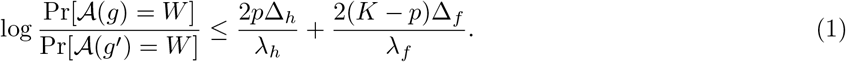

.Thus, if we set 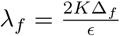 and 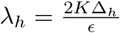, then (1) is equal to *ϵ* regardless of the value of *p*.

From the above and Theorem 1, Algorithm 3 satisfies *ϵ*-differential privacy.

## 4 Experiments and Discussion

We evaluated the usefulness of Algorithms 1 and 3 in terms of accuracy, rank error of the output, and run time. For the compressed sensing, we used Orthogonal Matching Pursuit [28]. In addition to our algorithms, we also experimented with the Laplace mechanism and the exponential mechanism using χ^2^-statistics as the score function and discuss the advantages of our methods. The Laplace mechanism and the exponential mechanism are shown in Algorithm 4 in Appendix A.1 and Algorithm 5 in Appendix A.2, respectively.

For accuracy, we varied the values of the privacy budget *ϵ* and the number of outputs *K* and calculated the accuracy rate of the top *K* data released upon using each mechanism. The value of *ϵ* was set from 0.01 to 1, because genomic data requires a high privacy guarantee. The value of *K* was set to values up to 10, because the goal of this study is to extract only significant SNPs. For the rank error, we evaluated the desirability of the output by measuring the error of the output data compared to their actual rank. Finally, we measured the run time of each mechanism by varying the input size. Note that in these experiments, the value of *η* in Algorithm 2 was set to 10 and the elements below 10 were dropped to 0, so that the information in the statistics vector was approximately represented as a sparse vector. In the future, we should develop a method to optimally determine the value of *η* under differential privacy, for example, by using the exponential mechanism.

In our experiments, we used simulation data for χ^2^-statistics in a χ^2^-test using a 3 × 2 contingency table. Assuming that the number of cases and controls is equal, we can use our algorithms because the *sensitivity* of the statistic is known to be 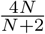, where *N* is the number of individuals [17].

### 4.1 Simulation Data

We generated data using two simulations, i.e, a small cohort with 100 individuals and 500 statistics, and a large cohort with 500 individuals and 5,000 statistics. Here, the χ^2^-statistic using a 3 × 2 contingency table is expressed using the values in Table 1 by the following equation: 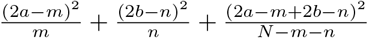.

**Table 1:**
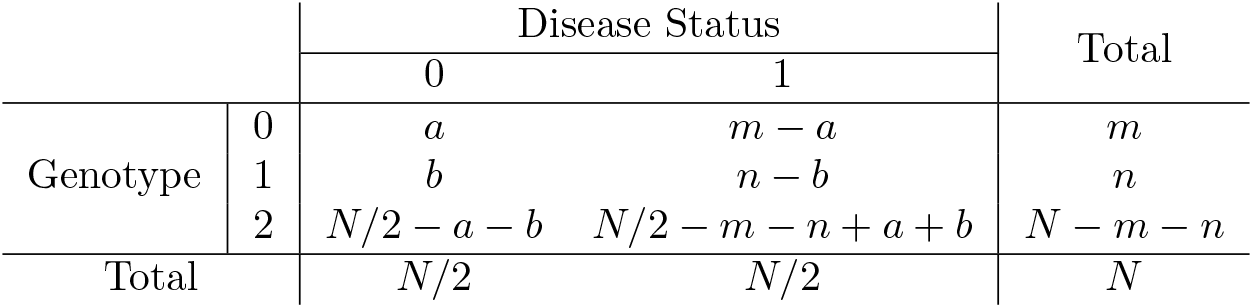
An example of 3 x 2 contingency table for χ^2^-test.

Let *N* = 100 for a small cohort and *N* = 500 for a large cohort, and we set *m, n, a,* and *b* using a binomial distribution with the following equations: 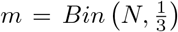, 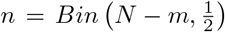, 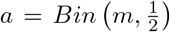, 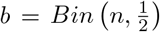. In addition, in order to generate 10 significant SNPs, the probabilities in the binomial distribution for calculating *a* and *b* were set as 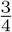 for a small cohort and 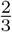 for a large cohort.

### 4.2 Results

#### 4.2.1 Accuracy

We varied the value of *ϵ* (from 0.01 to 1), and the value of *K* (1, 3, 5, and 10) and calculated the accuracy for the top *K* output using each mechanism. The results of the simulation data for a small cohort and a large cohort described in Subsection 4.1 are plotted in Figures 1 and 2.

**Figure 1:**
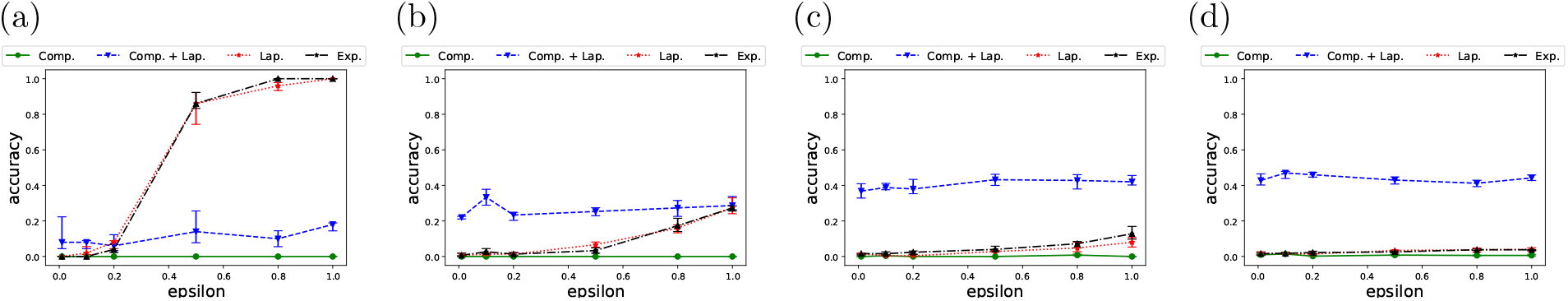
Accuracy of the *K* outputs when (a) *K* = 1, (b) *K* = 3, (c) *K* = 5, and (d) *K* = 10 in a small cohort. The green, blue, red, and black lines represent Comp., Comp. + Lap., Lap., and Exp., respectively.

**Figure 2:**
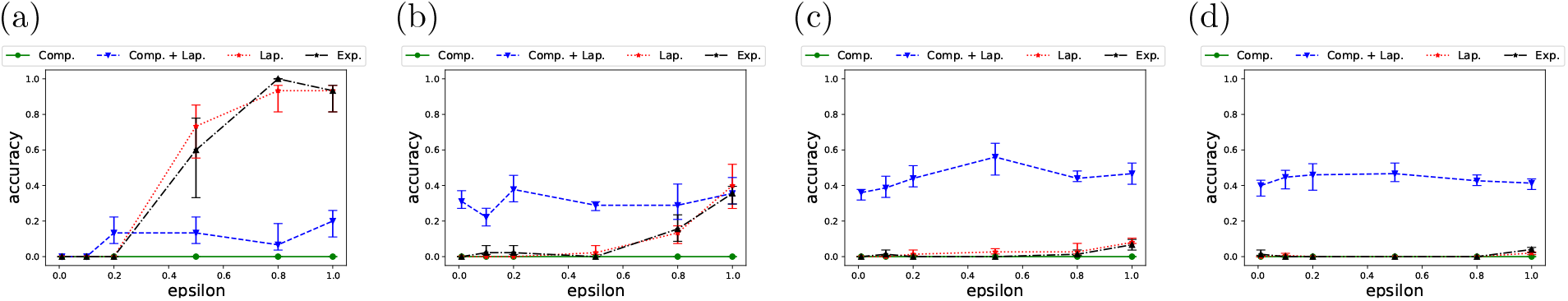
Accuracy of the *K* outputs when (a) *K* = 1, (b) *K* = 3, (c) *K* = 5, and (d) *K* = 10 in a large cohort. The green, blue, red, and black lines represent Comp., Comp. + Lap., Lap., and Exp., respectively.

Figure 1 shows that when *K* = 1, the Laplace and exponential mechanisms provide higher accuracy than our methods. However, the accuracy of these mechanisms decreases as *K* increases. The reason for this is that more noise we have to add to each statistic, the less we can distinguish between significant and non-significant SNPs. On the other hand, when *K* = 3, 5, and 10, our methods, which is a combination of the compressive and Laplace mechanisms, outperforms the other methods. More surprisingly, as *K* increases, higher accuracy is achieved, and when *K* = 10, approximately half of the output is correct. Even in the case of a large cohort, the outline in Figure 2 is like that in Figure 1, confirming that our mechanism achieved a higher accuracy than the others. However, our simple compressive mechanism did not provide an accurate output in any of the cases.

#### 4.2.2 Rank Error

We also evaluated the desirability of the output by calculating the rank error. The rank error represents the deviation from the actual rank and is expressed as 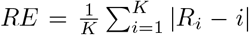, where *R_i_* is the true rank of the data output as the *i*-th. Even if the output does not exactly match the true data, when the rank error is small, it can be expected that the relatively high-rank data will be the output, which is desirable for practical use. The experimental results are shown in Figures 3 and 4, respectively.

**Figure 3:**
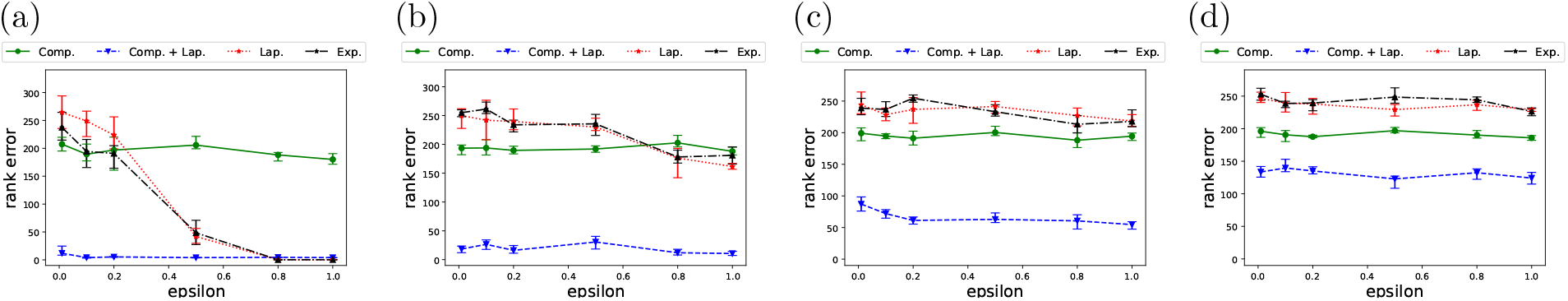
Rank error of the *K* outputs when (a) *K* = 1, (b) *K* = 3, (c) *K* = 5, and (d) *K* = 10 in a small cohort. The green, blue, red, and black lines represent Comp., Comp. + Lap., Lap., and Exp., respectively.

**Figure 4:**
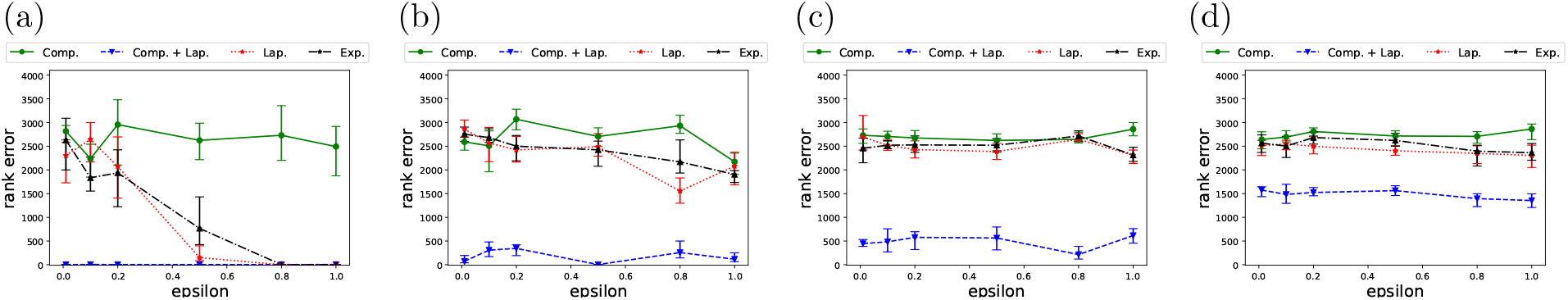
Rank error of the *K* outputs when (a) *K* = 1, (b) *K* = 3, (c) *K* = 5, and (d) *K* = 10 in a large cohort. The green, blue, red, and black lines represent Comp., Comp. + Lap., Lap., and Exp., respectively.

Figure 3 indicates that for a small cohort, our methods have much smaller output errors than existing mechanisms. For the Laplace and exponential mechanisms, as in Section 4.2.1, the utility decreased as *ϵ* decreased and as *K* increased. Our mechanisms did not significantly alter the rank error much, depending on the value ε. In particular, our method that combines the compressive and Laplace mechanisms performed better than the other methods in all cases. Even in the case of *K* =1, where the accuracy was not very high, the rank error was quite small, and we can expect a significant SNP output. In addition, the algorithm that involved only the compressive mechanism resulted in smaller rank errors than those that involved the Laplace and exponential mechanisms. However, in the case of a large cohort shown in Figure 4, a small rank error was not achieved upon using the compressive mechanism alone, instead a very high utility could be achieved upon using the combination with the Laplace mechanism.

In Appendix A.3, we show the results of additional experiments on accuracy and rank error for a larger cohort containing 25,000 SNPs. The results also indicate that our method outperforms the other mechanisms.

#### 4.2.3 Run Time

We varied the data size of the statistics data *g*, and measured the run time of each mechanism. Here, note that the parameters that affect the run time are only the dataset size |*g*|, i.e., the number of SNPs, and the number of outputs *K*. Thus, it does not depend on the number of individuals in the dataset or the value of ε. We conducted five runs for each case, and the average values are listed in Table 2.

**Table 2:** Run time [s] of each mechanism for each input size.

| $\vert g\vert $ | 500 | 1,000 | 2,500 | 5,000 | 10,000 | 25,000 |
| --- | --- | --- | --- | --- | --- | --- |
| Comp. | 6.52 | 28.37 | 567.2 | $9.2 \times 10^3$ | — | — |
| Comp. + Lap. | 2.96 | 10.03 | 136.6 | 588.2 | $2.1 \times 10^3$ | $7.9 \times 10^3$ |
| Lap. | $2.9 \times 10^{-4}$ | $3.9 \times 10^{-4}$ | $6.1 \times 10^{-4}$ | $1.1 \times 10^{-3}$ | $1.9 \times 10^{-3}$ | $5.6 \times 10^{-3}$ |
| Exp. | $1.6 \times 10^{-3}$ | $3.4 \times 10^{-3}$ | $8.7 \times 10^{-3}$ | $2.0 \times 10^{-2}$ | $3.2 \times 10^{-2}$ | $7.8 \times 10^{-2}$ |

With respect to the run time, our algorithms are inferior to the Laplace mechanism and the exponential mechanism owing to matrix multiplication and compressed sensing. Moreover, the run time of our method using only the compressive mechanism highly depends on the method to determine the number of nonzero entries. In this experiment, we considered elements larger than the threshold *η* in Algorithm 2 as nonzero entries, and the sparsity decreases with the increase in the data size. Consequently, the run time becomes longer. In contrast, the method that involved combining the compressive mechanism with the Laplace mechanism, which performed well in terms of accuracy and rank error, required only approximately 10 minutes to process 5, 000 data. However, when the data size was much larger than that shown in Tabke 2, e.g. when |*g*| = 100, 000, it took about two days. In the future, we will need to employ faster calculation methods in compressed sensing and more effective ways to determine the number of nonzero entries.

### 4.3 Real Data

We also evaluated the proposed mechanism using real data. The data we used were TDT statistics by Justice *et al.* [24]. Here, the *sensitivity* of TDT statistics for *N* families is 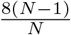 [32]. From their data, we took 2, 000 statistics and subjected them to our method of combining the compressive and Laplace mechanisms, Laplace mechanism, and the exponential mechanism to them. We also evaluated the exponential mechanism using the SHD score [32], which is applicable to datasets of the TDT statistics. Since there are six significant SNPs based on their statistics, we considered the cases where *K* = 1, 2, and 5. For each *K*, we calculated the probability of releasing those significant SNPs. The values of *ϵ* were set to 1, 2, and 5, which are larger than those used in the simulation studies, in order to also check the variation in the utility when using the Laplace and exponential mechanisms. The results are shown in Table 3.

**Table 3:**
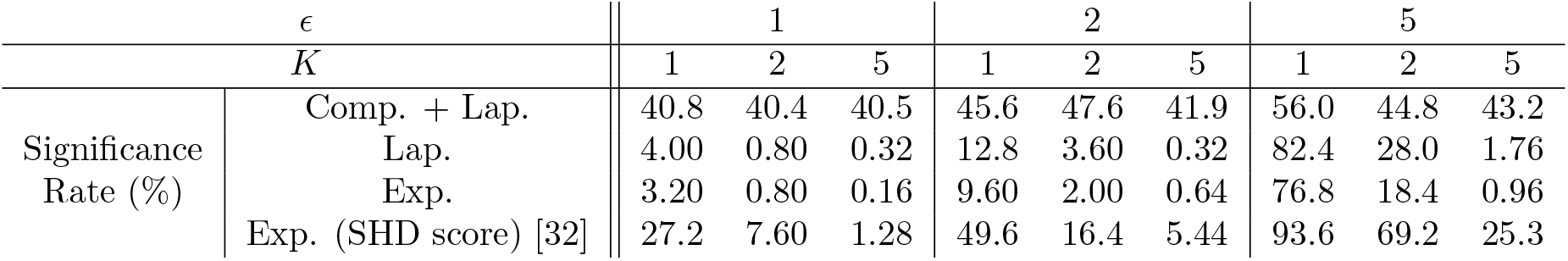
Probability of extracting significant SNPs.

The results indicate that our method is better than the Laplace and exponential mechanisms even for real data. We also found that our method can release significant SNPs with high probability even when *ϵ* becomes smaller and the number of outputs *K* increases. However, as with the results in Section 4.2, when *K* = 1, the Laplace and exponential mechanisms provide relatively high utility because the scale of adding noise is small compared to the difference among the original statistics. The exponential mechanism using the SHD score achieved relatively high accuracy, but was not as accurate as out method when the privacy level was high or when the number of outputs was large.

## 5 Conclusion

In this study, we proposed privacy-preserving methods using the compressive mechanism to release the top *K* significant SNPs based on a genomic dataset. Our first algorithm, which involves only the simple conmpressive mechanism, achieved smaller rank errors than existing algorithms involving the Laplace and exponential mechanisms in a small cohort. In our second method, we considered genomic data as a sparse vector and combined the compressive and Laplace mechanisms. This method outperformed the other methods in terms of accuracy as well as rank error by distinguishing significant SNPs from non-significant SNPs. We also theoretically proved that this method satisfies *ϵ*-differential privacy. In addition, we also obtained high performance in our experiments using real data, indicating that our method can ensure both high utility and privacy assurance.

One limitation of this study is that we could not achieve high accuracy using the compressive mechanism alone. We need to further improve the method by, for example, considering the position of nonzero elements in a sparse representation. In addition, for much larger datasets containing millions of SNPs, we would like to investigate additional methods to conduct calculation without using matrices and faster algorithms using the Fourier transform or other techniques. For further work, we need to consider wavelet transforms other than the Haar matrix and develop new score functions for the exponential mechanism.

## Acknowledgements

This work was supported by JSPS KAKENHI Grant 20H05967, 20K21827, and 21H05052. The supercomputing resource was provided by Human Genome Center, the Institute of Medical Science, the University of Tokyo.

## A Appendix

### A.1 Laplace Mechanism

An algorithm for releasing the top *K* significant SNPs based on genomic statistics data using the Laplace mechanism is shown in Algorithm 4.

**Algorithm 4.**
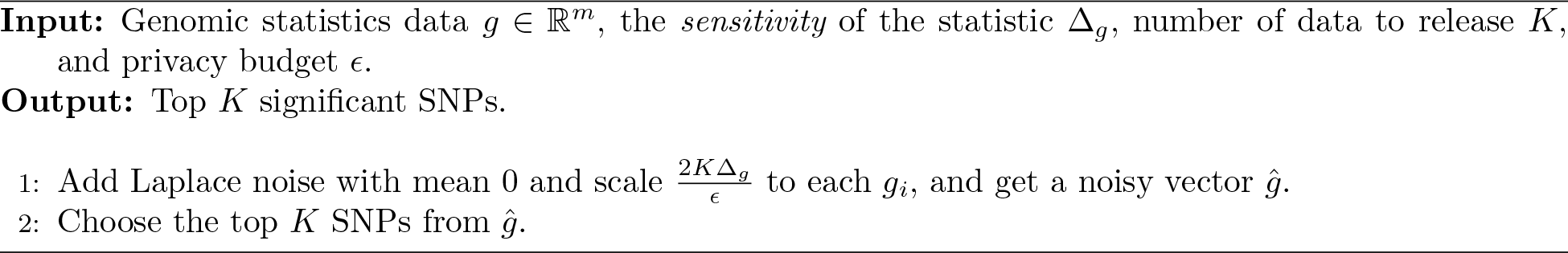
*ϵ*-differentially private algorithm for releasing the top *K* significant SNPs using the Laplace mechanism.

### A.2 Exponential Mechanism

An algorithm for releasing the top *K* significant SNPs based on genomic statistics data using the exponential mechanism is shown in Algorithm 5.

**Algorithm 5.**
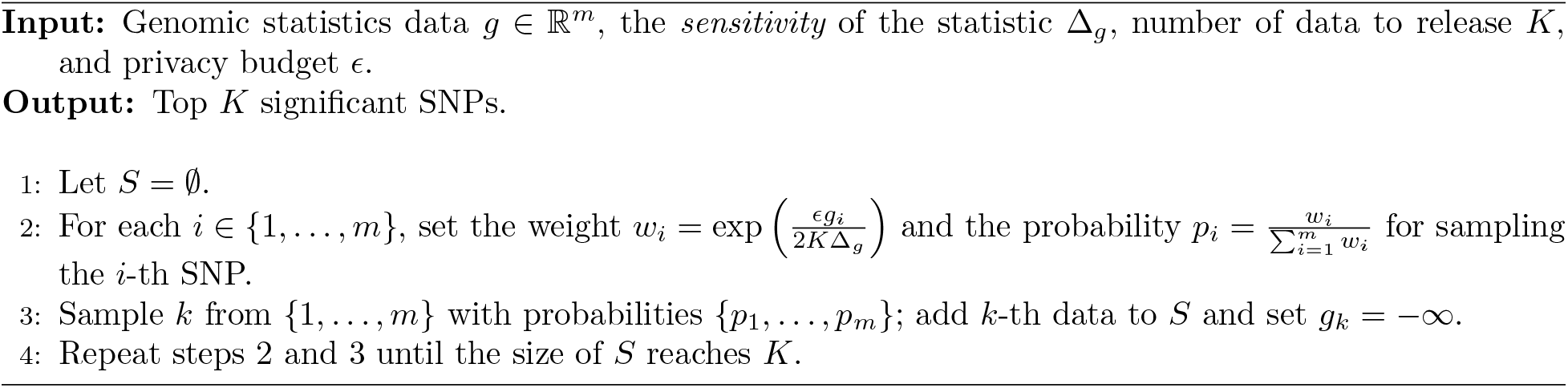
*ϵ*-differentially private algorithm for releasing the top *K* significant SNPs using the exponential mechanism.

### A.3 Additional Experiments

We also conducted experiments on accuracy and rank error for the case where the number of SNPs in the simulation dataset was 25, 000. The results are shown in Figures 5 and 6. When *K* =1, the existing Laplace and exponential mechanisms achieved high accuracy because the noise is small and the rank of the original statistics varied little. However, when *K* was increased or when *ϵ* was small, i.e., higher privacy was achieved, our combining method outperformed the other methods.

**Figure 5:**
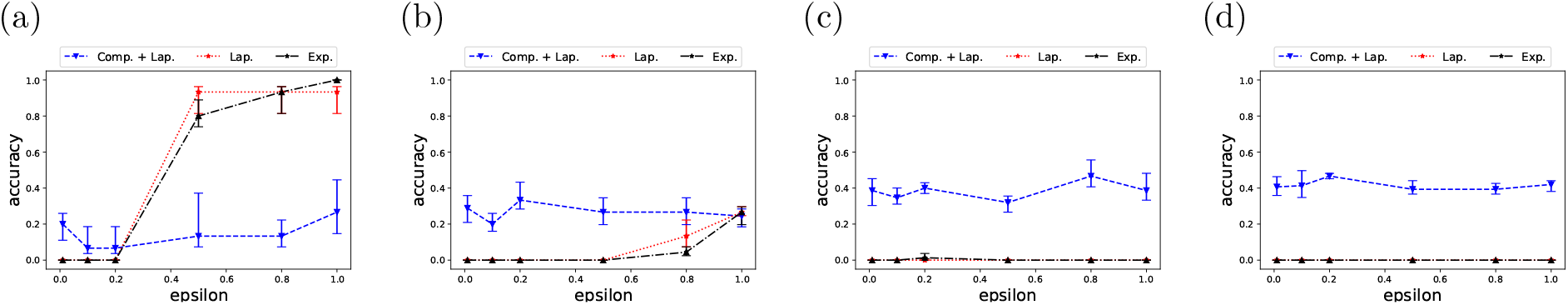
Accuracy of the *K* outputs when (a) *K* = 1, (b) *K* = 3, (c) *K* = 5, and (d) *K* = 10 in a large cohort with 25,000 SNPs. The blue, red, and black lines represent Comp. + Lap., Lap., and Exp., respectively.

**Figure 6:**
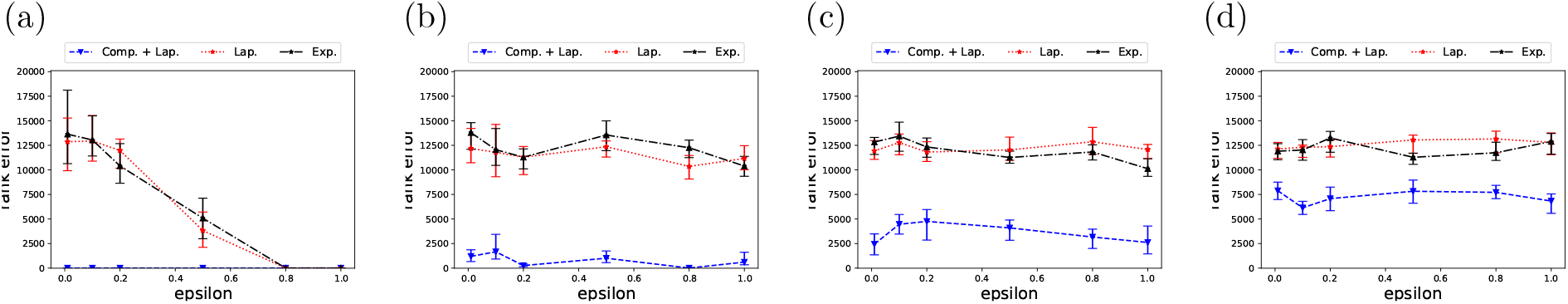
Rank error of the *K* outputs when (a) *K* = 1, (b) *K* = 3, (c) *K* = 5, and (d) *K* = 10 in a large cohort with 25,000 SNPs. The blue, red, and black lines represent Comp. + Lap., Lap., and Exp., respectively.

#### A.3.1 Accuracy

#### A.3.2 Rank Error

